# Bovine highly pathogenic avian influenza virus stability and inactivation in the milk byproduct lactose

**DOI:** 10.1101/2024.08.12.607665

**Authors:** Taeyong Kwon, Jordan T. Gebhardt, Eu Lim Lyoo, Mohammed Nooruzzaman, Natasha N. Gaudreault, Igor Morozov, Diego G. Diel, Juergen A. Richt

## Abstract

A bovine isolate of highly pathogenic avian influenza H5N1 virus was stable for 14 days in a concentrated lactose solution at under refrigerated conditions. Heat or citric acid treatments successfully inactivated viruses in lactose. This study highlights the persistence of HPAIV in lactose and its efficient inactivation under industrial standards.

---

Recently, clade 2.3.4.4b viruses were detected in dairy cattle populations in the United States (1, 2). Infected cattle exhibited reduced appetite, fever, mild respiratory symptoms, reduced milk production, and changes in milk quality (3). High levels of virus shedding in milk have been detected in affected cows, with virus titers ranging from 10^4^ to 10^8.8^ TCID_50_/mL (3). It is presumed that a single introduction from wild birds to cattle occurred and subsequently the movement of subclinical cattle played a significant role in the spread to multiple sites (4). Four human infections with HPAI H5N1 viruses following exposure to dairy cattle have been reported so far (5, 6). Given the high titer of virus in milk and the potential for H5N1 transmission via raw milk and its byproducts to humans and agricultural animals, it is essential to develop appropriate processes to inactivate the bovine H5N1 virus in these substrates to mitigate the risk of transmission. The current knowledge and techniques have focused on pasteurization of milk, which is widespread within the dairy industry and has been shown to be effective for HPAI viruses (7-9). Consumption of contaminated materials is presumably considered a major route of HPAI infection in pet and wild mammals (10). Therefore, additional research is needed to validate inactivation processes in milk and its byproducts, such as dried whey, whey permeate, and lactose, which are used for animal nutrition in agriculture. Therefore, this study aimed to determine the stability of a bovine H5N1 isolate in the milk byproduct lactose and to evaluate two inactivation methods using industrial procedures.

The bovine isolate of HPAI H5N1 clade 2.3.4.4b, isolate A/Cattle/Texas/063224-24-1/2024 (3), was propagated and titrated MDCK cells. The virus stock was mixed with a concentrated lactose solution at 1:10 dilution, and 1 mL of contaminated lactose were incubated at refrigerated temperatures and titrated on MDCK cells.

For inactivation, virus-spiked lactose at a 1:10 dilution was subjected to heat or citric acid treatments. A total of 1 mL of the H5N1-spiked lactose was incubated at 63, 66, or 99 °C for up to 30 min and cooled down on ice water for at least 30 minutes. The temp 63 °C for 30 min was chosen since it is used by the food industry for pasteurization. Other temperatures were chosen to evaluate the efficacy of elevated temperatures and shorter time. For citric acid treatment, 1 mL of the H5N1 spiked lactose was mixed with 1 mL of different levels of citric acid and incubated at a refrigerated temperature. After a defined incubation time, the sample was neutralized by adding 1N NaOH. All samples were titrated on MDCK cells, and the presence of infectious viruses were visualized by immunofluorescence assay.

The H5N1 virus was rather stable in lactose at a refrigerated temperature, and there was only a 1-log reduction after incubation for 14 days (Figure 1). Heat treatment at 63 °C for 5 minutes resulted in 3-log reduction in both high and low dose samples, and all samples were virus negative at 15 and 30 minutes (Table 1). No infectious virus was isolated after heat treatment at 66 °C and 99 °C for a minimum of 5 min. Next, we investigated the effect of citric acid treatment on virus inactivation in the concentrated lactose solution. In high dose samples, low levels of virus were still present after treatment with 0.2% and 0.4% citric acid for up to 1 hour, but virus was inactivated after 0.6% treatment. For low dose samples, 0.2% and 0.4% citric acid treatment successfully inactivated H5N1 within 5 minutes, and 0.1% treatment was effective at inactivating viruses after 15 minutes contact time (Table 1).

**Figure 1.**
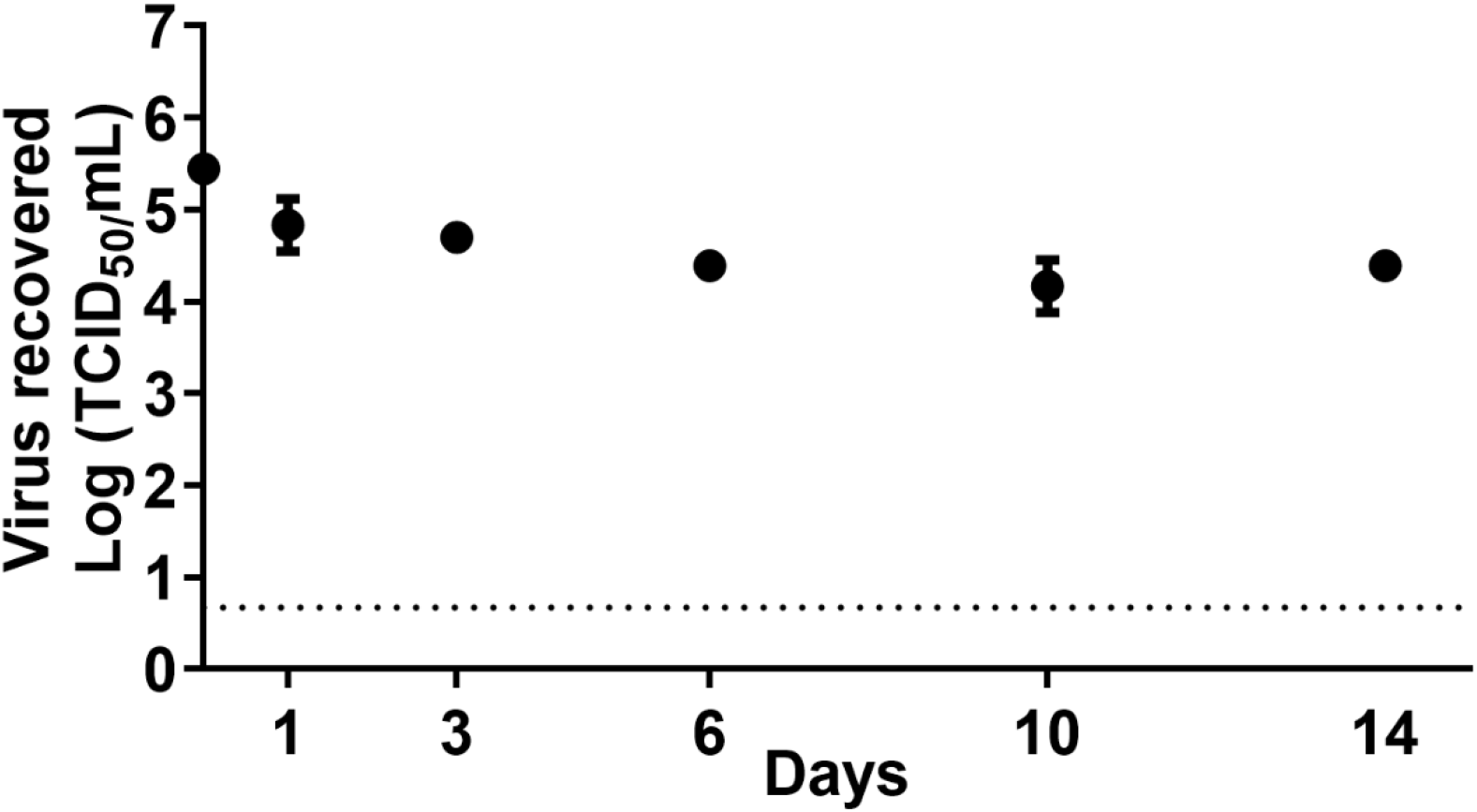
Stability of the bovine isolate of HPAI H5N1 clade 2.3.4.4b in a concentrated lactose solution. The virus was mixed with whole milk and lactose at 1:10 dilution and incubated at a refrigerated temperature. At each time point, the samples were titrated on MDCK cells.

**Table 1.** Inactivation of the bovine isolate of HPAI H5N1 clade 2.3.4.4b.

| Treatment | Virus dose | Condition | Time and virus titer <sup>1</sup> (TCID<sub>50</sub>/mL) |  |  |  |
| --- | --- | --- | --- | --- | --- | --- |
|  |  |  | 0 min | 5 min | 15 min | 30 min |
| Heat | High | 63 °C | 3.59E+05 | 4.64E+02 | Negative | Negative |
|  |  | 66 °C |  | Negative | Negative | Negative |
|  |  | 99 °C |  | Negative | Negative | Negative |
|  | Low | 63 °C | 1.29E+03 | 3.78E+00 | Negative | Negative |
|  |  | Acidulant level | 0 min | 20 min | 40 min | 60 min |
| Citric acid<sup>2</sup> | High | 0.2% | 2.15E+05 | 7.74E+00 | 5.99E+00 | 6.14E+00 |
|  |  | 0.4% |  | 3.78E+00 | 3.68E+00 | 4.64E+00 |
|  |  | 0.6% |  | Negative | Negative | Negative |
|  | Low | Untreated | 1.29E+03 | 1.00E+03 | 1.29E+03 | 1.29E+03 |
|  |  | 0.1% |  | 3.68E+00 | Negative | Negative |
|  |  | 0.2% |  | Negative | Negative | Negative |
|  |  | 0.4% |  | Negative | Negative | Negative |

<sup>1</sup> The limit of detection of this assay was 4.64 TCID<sub>50</sub>/mL.

Our findings highlight H5N1 virus survival in the concentrated lactose solution for up to 14 days at holding temperature and emphasize the successful inactivation by pasteurization (63 °C) and citric acid treatment. In this study, samples with high virus titers are consistent with the observed range of virus titers in milk from infected cows (3), and the low titer samples would theoretically be possible if milk from infected animals is diluted with milk from non-infected animals. Thus, the viral titers used in the current experiment are representative of possible virus levels in milk and its coproducts.

This is the first study to investigate the inactivation of bovine HPAI H5N1 virus in the coproduct concentrated lactose solution which is frequently used as a feed ingredient in agricultural animals including pigs as well as for other purposes. In summary, H5N1 contaminated milk byproducts might pose a risk to animal health if consumed untreated. This study provides insights on the persistence of HPAIV in dairy byproducts and effective inactivation strategies under industrial standards.

## Acknowledgments

This study was supported through the National Bio and Agro-Defense Facility (NBAF) Transition Fund from the State of Kansas, and the MCB Core of the Center on Emerging and Zoonotic Infectious Diseases (CEZID) of the National Institutes of General Medical Sciences under award number P20GM130448, and the NIAID supported Center of Excellence for Influenza Research and Response (CEIRR) under contract number 75N93021C00016.

## Biographical Sketch

Dr. Kwon is a PhD candidate at Kansas State University, Manhattan, Kansas, USA. His research interests include transboundary animal diseases and emerging zoonotic diseases.

## Address for Correspondence

Juergen A. Richt, Kansas State University, 1800 Denison Avenue, Manhattan, KS 66506, USA

## References

1. Animal and Plant Health Inspection Service. Detections of Highly Pathogenic Avian Influenza (HPAI) in Livestock. 2024.

2. Singh G, Trujillo JD, McDowell CD, Matias-Ferreyra F, Kafle S, Kwon T, et al. Detection and characterization of H5N1 HPAIV in environmental samples from a dairy farm. 2024.

3. Caserta LC, Frye EA, Butt SL, Laverack M, Nooruzzaman M, Covaleda LM, et al. Spillover of highly pathogenic avian influenza H5N1 virus to dairy cattle. Nature. 2024 Jul 25.

4. Nguyen T-Q, Hutter C, Markin A, Thomas M, Lantz K, Killian ML, et al. Emergence and interstate spread of highly pathogenic avian influenza A(H5N1) in dairy cattle. bioRxiv. 2024.

5. Uyeki TM, Milton S, Abdul Hamid C, Reinoso Webb C, Presley SM, Shetty V, et al. Highly Pathogenic Avian Influenza A(H5N1) Virus Infection in a Dairy Farm Worker. N Engl J Med. 2024 Jun 6;390(21):2028–9.

6. Centers for Disease Control and Prevention. CDC Reports Fourth Human Case of H5 Bird Flu Tied to Dairy Cow Outbreak. 2024.

7. Kaiser F, Morris DH, Wickenhagen A, Mukesh R, Gallogly S, Yinda KC, et al. Inactivation of Avian Influenza A(H5N1) Virus in Raw Milk at 63 degrees C and 72 degrees C. N Engl J Med. 2024 Jun 14.

8. Guan L, Eisfeld AJ, Pattinson D, Gu C, Biswas A, Maemura T, et al. Cow’s Milk Containing Avian Influenza A(H5N1) Virus - Heat Inactivation and Infectivity in Mice. N Engl J Med. 2024 May 24.

9. Spackman E, Jones DR, McCoig AM, Colonius TJ, Goraichuk I, Suarez DL. Characterization of highly pathogenic avian influenza virus in retail dairy products in the US. MedRxiv. 2024.

10. Adlhoch C, Fusaro A, Gonzales JL, Kuiken T, Melidou A, Mirinaviciute G, et al. Avian influenza overview April - June 2023. EFSA J. 2023 Jul;21(7):e08191.

